# CeutaOPEN: Individual-based field observations of breeding snowy plovers *Charadrius nivosus*

**DOI:** 10.1101/2019.12.20.884825

**Authors:** Luke J. Eberhart-Phillips, Medardo Cruz-López, Lydia Lozano-Angulo, Salvador Gómez del Ángel, Wendoly Rojas-Abreu, Clemens Küpper

## Abstract

Shorebirds (*partim* members of order Charadriiformes) have a global distribution and exhibit remarkable variation in ecological and behavioural traits that are pertinent to many core questions in the fields of evolutionary ecology and conservation biology. Shorebirds are also relatively convenient to study in the wild as they are ground nesting and often occupy open habitats that are tractable to monitor. Here we present a database documenting the reproductive ecology of 1,600 individually marked snowy plovers (*Charadrius nivosus*) monitored between 2006 and 2016 at Bahía de Ceuta (23°54 N, 106°57 W) – an important breeding site in north-western Mexico. The database encompasses various morphological, behavioural, and fitness-related traits of males and females along with spatial and temporal population dynamics. This open resource will serve as an important data repository for addressing overarching questions in avian ecology and wetland conservation during an era of big data and global collaborative science.

## Background & Summary

Longitudinal data on individuals living in the wild represent the gold standard for research in organismal ecology, as subjects are sampled repeatedly over multiple stages of their life-history while being exposed to the natural evolutionary pressures of their native environments^1^. These types of data have offered evolutionary ecologists valuable insights into the selective processes that effect species over multiple generations such as, for example, the role of stochastic climate events shaping the beak morphologies of Darwin’s Finches^2^, the predator-prey cycles of mammal communities on the Serengeti^3^, or the demographic dynamics of alpine plants^4^ and animals^5^ in response to climate change. However, collecting field data over many consecutive years while following standardized methods requires substantial labour and consistent funding. Due to these challenges, raw longitudinal field data from wild populations are rarely made open to the public^6^ – thus limiting the transparency and reproducibility of published research methods and results in evolutionary ecology. Furthermore, releasing raw data has the potential benefit of stimulating more substantive discussion and criticism within the scientific community, which can advance research topics and forge productive collaborations. Here, we offer an open access database of our raw field observations over an 11-year period of 1,600 uniquely marked individuals from an important breeding population of snowy plovers (*Charadrius nivosus*) in Mexico.

*Charadrius* plovers are small ground-nesting shorebirds that occur worldwide. As a group, plovers present a model system for investigating fundamental and applied topics in organismal biology as they occupy open habitats that are easy to monitor and experimentally manipulate, and they exhibit intra- and interspecific variation in several behavioural, ecological, and demographic traits. For example, plovers display remarkable diversity and plasticity in breeding tactics with sex roles during courtship, mating, and parental care varying appreciably among populations both between and within species^7^. The snowy plover is native to North America^8^ and is one of the least abundant shorebirds on the continent (estimated population size: 25,869) with many populations in decline and requiring intensive management^9^. Apart from being a public icon of avian conservation, snowy plovers have also increasingly captured the spotlight for their intriguing ecology and life-history. Their unusual biology features a rare breeding behaviour characterized by highly dispersive polyandry and male-biased uniparental care^10,11^.

In this data descriptor we present CeutaOPEN – an open-access database containing the raw data from our fieldwork between 2006 and 2016 monitoring a breeding population of snowy plovers at Bahía de Ceuta, a subtropical lagoon on the coastal plain of north-western Mexico (23°54′ N, 106°57′ W). The database includes individual-based observations of reproductive effort, movements, morphometrics, and social behaviour (Fig. 1). Previously, we have used subsets of these data to report on a wide variety of topics in organismal biology, including sex ratio variation^12^, population viability^13^, courtship behaviour^14^, incubation behaviour^15^, parental care^16^, ontogeny^17^, chronobiology^18^, camouflage mechanisms^19^, offspring desertion^20^ and mating system dynamics^21^. The motivation for making our database open is to provide evolutionary ecologists with an accessible resource that will serve as an important repository for addressing overarching questions in organismal biology and conservation. Here we describe our field methods for collecting the observations presented in the database, we summarize the contents of the database, and we provide a code-based tutorial demonstrating how to import and query the database within the R environment and conduct, for example, a simple analytical workflow to investigate sex-specific ontogeny.

**Figure 1.**
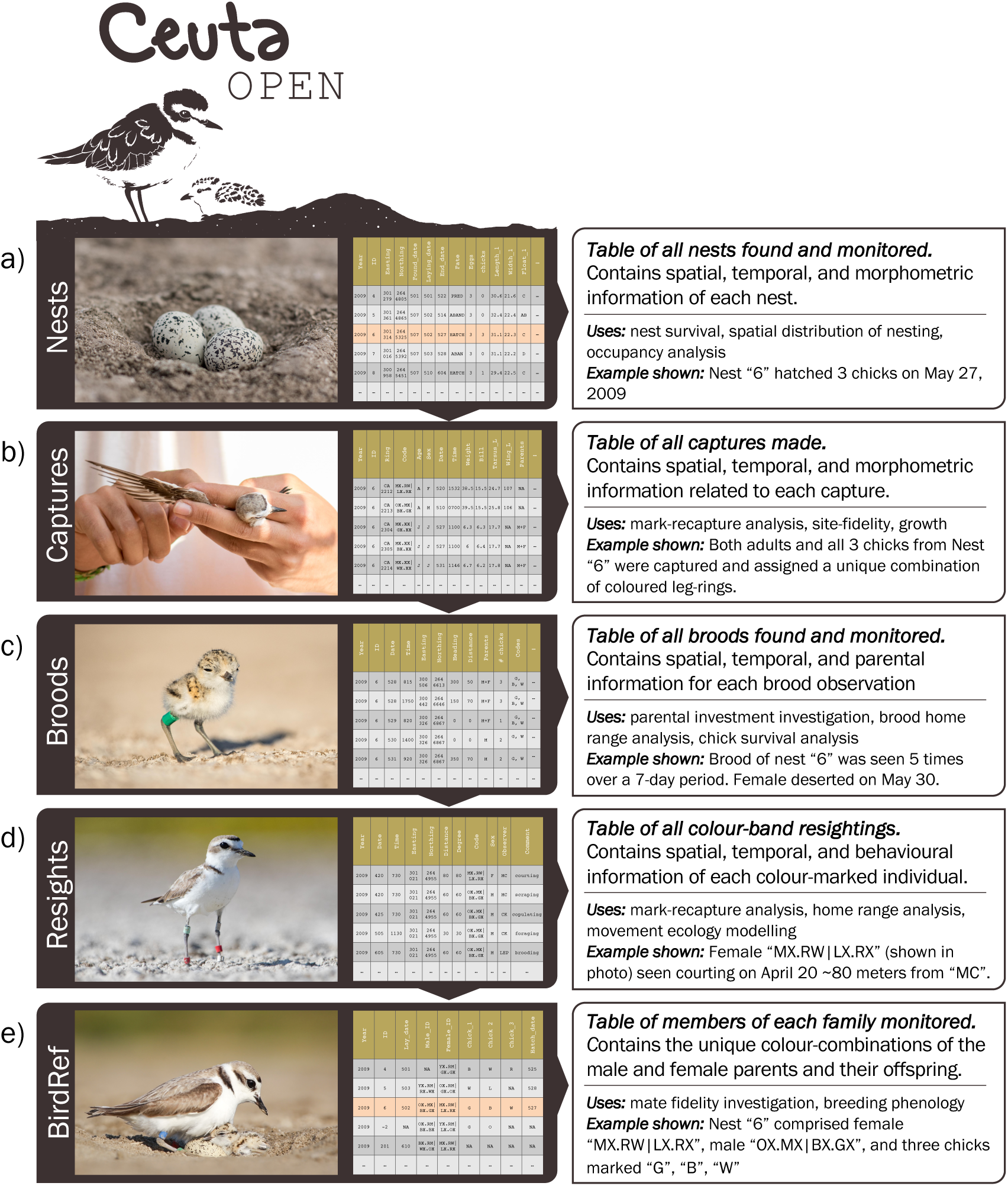
Schematic of the CeutaOPEN database. During fieldwork, data collection was divided across five main tasks: a) nest monitoring, b) captures of adults and chicks, c) brood monitoring, d) resightings of individually colour ringed adults, and e) determining the identity of all breeding pairs and their offspring. The data obtained during each of these activities are structured in our database as the five tables shown here that contain a common variable such as a nest “ID” or a bird “code”, that can be utilized by the user for relational queries.

## Methods

### Study area

Plovers breeding in Bahía de Ceuta mainly concentrate their activities on 200 ha of salt flats that contain several abandoned evaporation ponds. This habitat (hereafter “salina”) is surrounded by red mangrove (*Rhizophora mangle*) and characterized by sparse vegetation and open substrates. Nesting typically commences in late March or April when flood waters recede and concludes by mid-July when rains and high tides submerge the salina again. Our monitoring effort throughout the 11-year study period was focused on the largest contiguous section of salt flats in the study area where the vast majority of breeding activity occurs (Fig. 2a, b). However, in drought years or at the end of the breeding season when tidewaters had retreated, we made observations of plovers nesting and tending broods in several small pockets of salina adjacent to the main study site (Fig. 2a).

**Figure 2.**
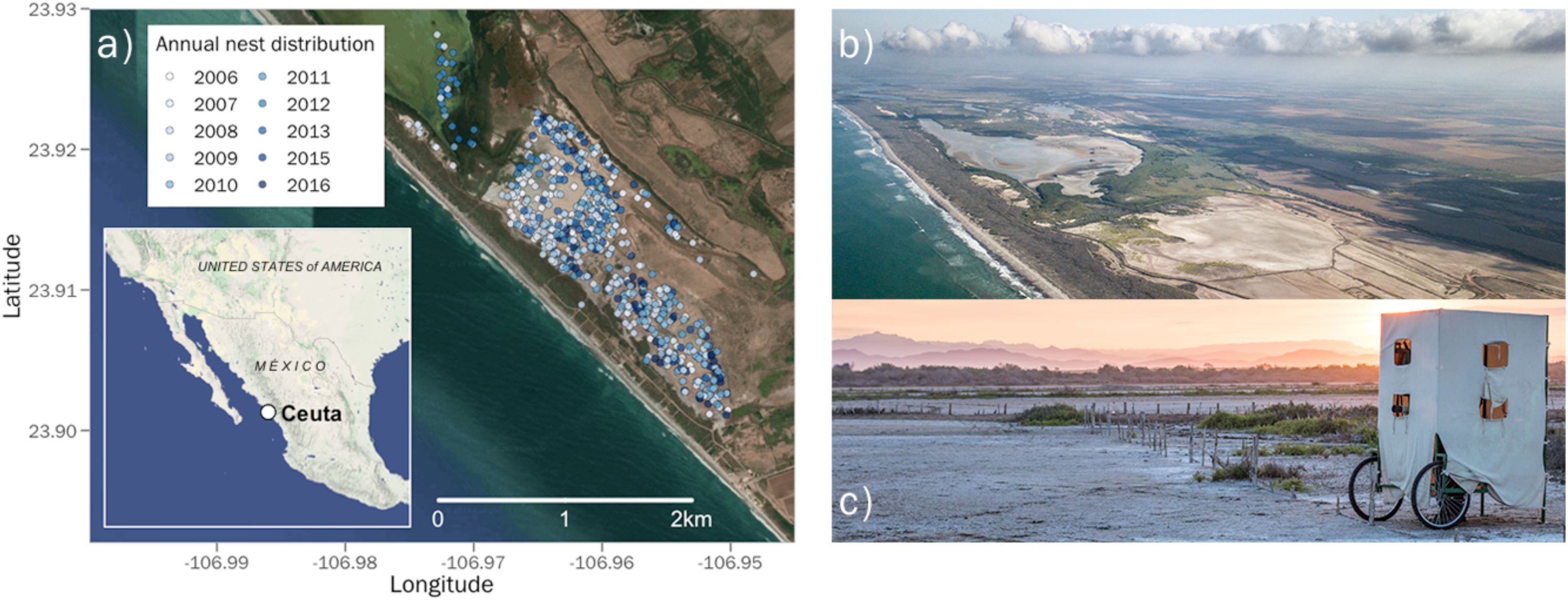
a) Map of the Bahía de Ceuta study site and photos of the b) salina breeding habitat and c) a mobile hide in which observers conduct non-invasive field work.

### Data collection

Over the 11-year study period, we monitored the population daily between April and July. We used a car and mobile hides^22^ (Fig. 2c) to search for nests, broods, and determine the identity breeding plovers with binoculars and scopes. During fieldwork, our data collection was divided across four main field tasks: 1) nest monitoring, 2) captures of adults and chicks, 3) brood monitoring, and 4) resightings of individually colour ringed adults. The data obtained during each of these activities are structured in our database as tables (Fig. 1) containing a common variable such as a nest “ID” or a bird “code”, that can be utilized by the user for relational queries. The format of these tables was originally conceived by Tamás Székely^23^. Fieldwork permits to collect the data presented in CeutaOPEN were granted by the Secretaría de Medio Ambiente y Recursos Naturales (SEMARNAT). All of our field activities were performed in accordance with the approved ethical guidelines outlined by SEMARNAT. Here we explain the details of our data collection pertinent to the database.

#### Nest Data

We regularly searched for nests (Fig. 1a) and incubating plovers by traversing the salina on foot, by car or in a mobile hide^19^(Fig. 2c). Upon discovery, we recorded the nest’s geographic location, the found date and time, and measured the width and length of each egg in the clutch with callipers. To estimate the initiation-date of the clutch (i.e., date when the first egg was laid), we floated each egg in a jar of water and scored the embryonic stage of development according to a calibrated table^24^. For hatched clutches that were initially discovered more than 10 days after laying, we estimated initiation-date by subtracting 25 days (i.e., the mean incubation time in our population^17^) from the hatching date and subtracting an additional 5 days to account for a 2-day egg laying interval^11^. We checked nests every 2–7 days to assess survival and identify tending parents.

#### Capture Data

We captured plover chicks by hand and adults using funnel traps on broods or nests. To individually identify members of the population, we assigned adults a unique combination of three to four colour leg rings and an alpha-numeric metal ring (Fig. 1b). Likewise, we marked chicks less than 2 weeks old with a single colour ring and a metal ring (Fig. 1c). Given our intensive nest search and capture efforts, we are confident that we ringed the vast majority of chicks (>95%) and breeding adults (>85%) in the local breeding population every year. During captures, we sampled the metatarsal vein of chicks or the brachial vein of adults and drew ∼25–50 μL of blood for subsequent genetic analyses. Additionally, we measured body mass, bill length, tarsus length, and wing length for all captured individuals (Fig. 3). As snowy plovers exhibit only minor sexual dimorphisim in plumage and body size, we molecularly determined sex using the Z-002B marker^25^ and verification with the Calex-31 marker located on the W chromosome^26^ for all adults and chicks captured before 2014. For PCR conditions see ref. 17.

**Figure 3.**
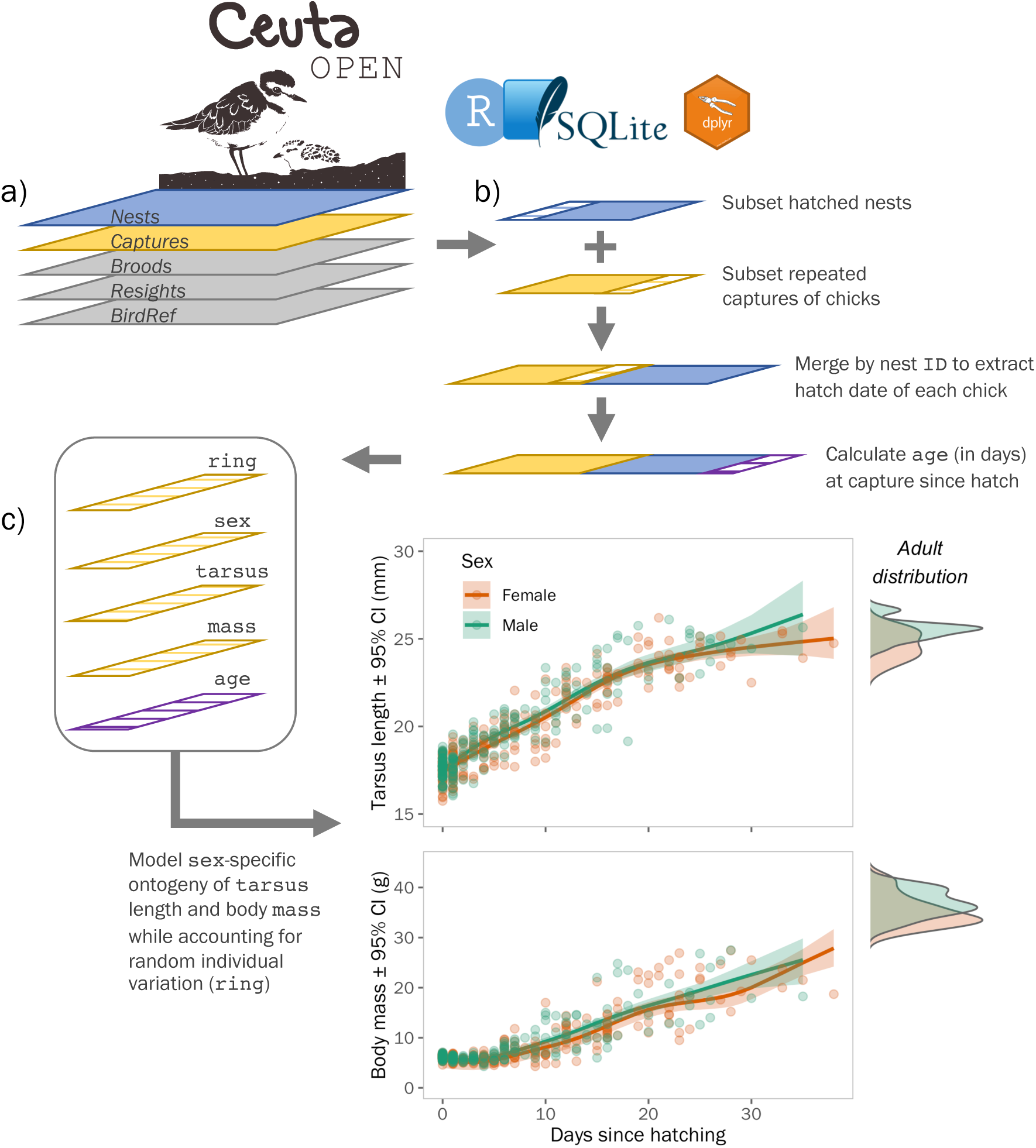
Schematic of an example analytical workflow using CeutaOPEN within the R environment to investigate how chick morphometric data may be used to study growth and ontogeny. a) Import the database into R using the RSQLite package. b) Use the dplyr package to join the ‘*Captures’* table with the ‘*Nests’* table by nest “ID” to determine the “hatch_date” of each captured individual. Subset the result to individuals that have repeated captures and calculate the “age” at capture by subtracting the “hatch_date” by the capture “date”. c) Use the bamlss^32^ package (i.e., “Bayesian Additive Models for Location, Scale, and Shape”) to determine the sex-specific growth trends while controlling for repeated measures within individuals and random annual variation. Plot the fitted values to visualise the trends (see Supplementary File for more details).

#### Brood Data

Similar to our data collection of nests, we resighted broods every 1–7 days to assess chick survival and determine sex-specific patterns of parental care and desertion. Each brood observation includes the time, distance and azimuth to the brood, geographic location of the observer, number of chicks seen, and the identity of them and their parents.

#### Resight Data

We typically resighted colour ringed individuals opportunistically whilst in the field. Since 2009 we surveyed the entire salina within a single day at least once during the breeding season to record all colour ringed individuals present (Fig. 1d). Like with our brood data, each resight includes the distance and azimuth to the individual, the geographic location of the observer, and any noteworthy comments pertaining to the individual’s behaviour.

## Code availability

To assist users with accessing and querying our database, we have written an accompanying RMarkdown document (Supplementary File 1) that provides a commented workflow for utilizing CetuaOPEN with the RSQLite^27^ and dplyr^28^ packages in R.

## Data Records

Our database and all other files described in this manuscript are stored in a publicly available OSF repository^29^. The file Ceuta_OPEN_v1.sqlite contains the SQL (Structured Query Language) database of four tables containing our raw observations collected during routine fieldwork (Nests, Captures, Broods, and Resights), and a fifth table (BirdRef) that uses relational information to summarize the identities of the parents and offspring belonging to each nest and subsequent brood. The structure of these tables is defined in Tables 1, 2, 3, 4, and 5 below. This Data Descriptor is based on version 1.0 of the CeutaOPEN database.

**Table 1.**
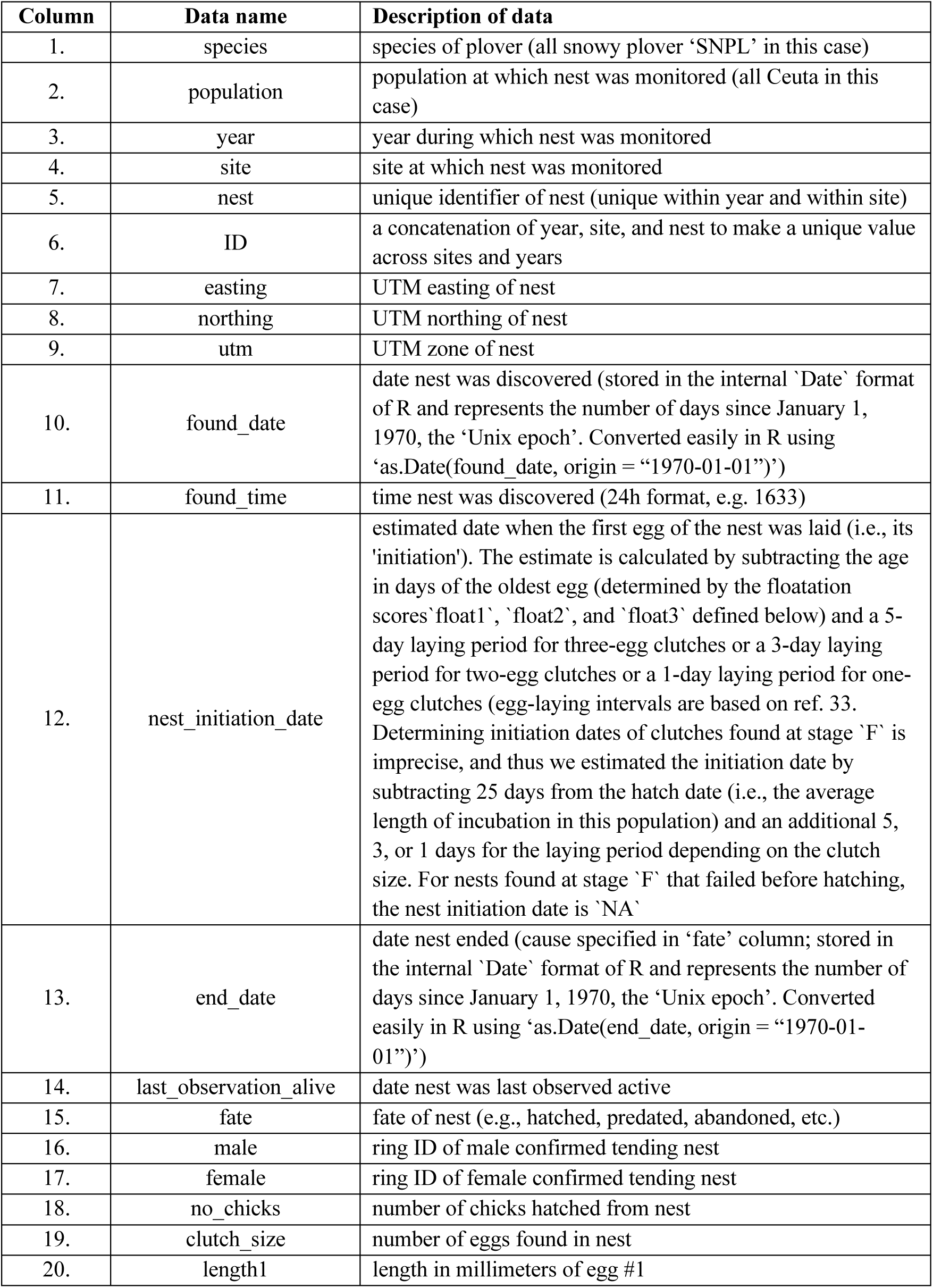
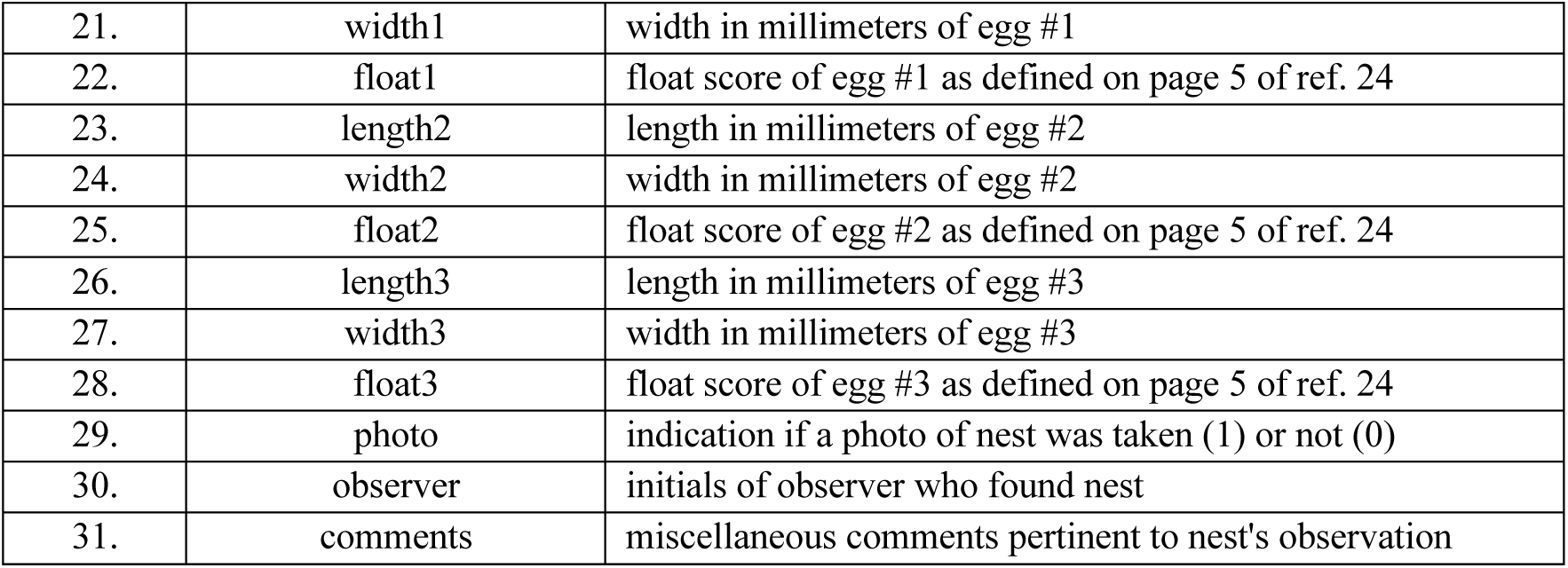
Nest data of snowy plovers breeding in Bahía de Ceuta, Mexico, between 2006 and 2016. This dataset contains information on egg dimensions, laying phenology, nest fate, geographic location, and the identity of incubating parents. These data can be used to assess individual reproductive effort and success, mate and site fidelity, and senescence, for example.

**Table 2.** Capture data of snowy plovers breeding in Bahía de Ceuta, Mexico, between 2006 and 2016. This dataset contains information on bird morphology, age, sex, capture time and location, and the identity of the individual. These data can be used to assess apparent survival with mark-recapture models, site fidelity, and growth rates of chicks, for example.

| Column | Data name | Description of data |
| --- | --- | --- |
| 1. | species | species of plover (all snowy plover ‘SNPL’ in this case) |
| 2. | population | population at which capture was made (all Ceuta in this case) |
| 3. | year | year during which capture was made |
| 4. | site | site at which capture was made |
| 5. | nest | unique identifier of nest at which capture was made (unique within year and within site). If capture was made at a brood originating from an unknown nest, the ID is negative (e.g., -2). |
| 6. | ID | a concatenation of year, site, and nest to make a unique nest ID |
| 7. | ring | alpha-numeric code of metal ring assigned to captured individual |
| 8. | code | color-ring combination assigned to captured individual. The scheme can be noted as XX.XX\|XX.XX where X indicates a possible position for a color (or metal) ring, the full-stop marks the position of the 'knee-joint' and the pipe divides the left and right leg. Thus the readout is "left above . left below \| right above . right below". See page 9 of ref. 24 for more details. |
| 9. | age | age of captured individual (J = juvenile (chicks and first-years), A = adult (second years and older)) |
| 10. | field_sex | sex of individual determined in the field based on ornamentation and other clues (e.g., time of capture, parental care, etc.), where F = female, M = males, and J = unknown sexed juvenile |
| 11. | mol_sex | sex of individual determined in the lab with the P2/P8 and Calex-31 markers (for our PCR conditions see ref. 17), where F = female, M = males, U = insufficient molecular evidence (e.g., markers failed), and NA = individual not molecularly sex-typed. Note: all birds initially captured in years after 2013 have not yet been molecularly sex-typed |
| 12. | sex | sex of captured individual (F = female, M = males, J = unknown sexed juvenile) |
| 13. | easting | UTM easting of capture |
| 14. | northing | UTM northing of capture |
| 15. | utm | UTM zone of capture |
| 16. | date | date capture was made (stored in the internal ‘Date’ format of R and represents the number of days since January 1, 1970, the ‘Unix epoch’. Converted easily in R using ‘as.Date(date, origin = “1970-01-01”)’) |
| 17. | time | time capture was made (24h format, e.g. 1633) |
| 18. | parents | parents attending captured individual (if age = "J") at time of observation (0 = no parent present; 1 = one parent (not identified whether male or female); 2 = female only (2+ when female certainly identified, whilst male uncertain); 3 = male only (3+ when male certain, whilst female uncertain, i.e., opposite of 2+); 4 = both present) |
| 19. | weight | weight in grams of captured individual |
| 20. | bill | length in millimeters of upper mandible of captured individual. Measured as the distance between the tip of the forehead feathering at the base of the upper bill, along the ridge of the culmen, and the tip of the bill (also known as the "exposed culmen" measurement; *sensu* page 8 of ref. 34 |
| 21. | left_tarsus | length in millimeters of left tarsus of captured individual. Measured as the distance between the notch at the end of the lateral condyle of the tibiotarsus on the backside of the leg, to the last tarsal scute on the front of the leg at the base of the foot (also known as the "outside tarsus" or "diagonal tarsus" measurement; *sensu* page 11 of ref. 34 |
| 22. | right_tarsus | same as left_tarsus measurement above but for right leg of captured individual |
| 23. | left_wing | length in millimeters of left wing of captured individual. Measured as the distance from the carpal joint (the bend of the wing) to the longest primary feather whilst flattening the wing and straightening the primaries (also known as the "maximum flat" or "flattened and straightened" measurement; *sensu* page 6 of ref. 34 |
| 24. | right_wing | same as left_wing measurement above but for right wing of captured individual |
| 25. | blood | indication if blood from captured individual was collected (1) or not (0) |
| 26. | moult | primary molt score of captured individual. Scored as the stage of the moult and the number of feathers at that stage. See ref. 35 for more details. |
| 27. | fat | fat score of captured individual, scored as the amount of visible fat in the furcular region or tracheal pit. See ref. 35 for more details. |
| 28. | lice | indication if feather lice from captured individual were collected (1) or not (0) |
| 29. | faecal | indication if faeces from captured individual was collected (1) or not (0) |
| 30. | photo | indication if a photo of captured individual was taken (1) or not (0) |
| 31. | observer | initials of observer making capture |
| 32. | comments | miscellaneous comments pertinent to capture event |

**Table 3.** Brood data of snowy plovers breeding in Bahía de Ceuta, Mexico, between 2006 and 2016. This dataset contains information on the time and location of a brood observation, the identity and number of chicks seen alive, and the identity of the parents tending chicks. These data can be used to assess parental investment, brood home range, and chick survival, for example.

| Column | Data name | Description of data |
| --- | --- | --- |
| 1. | species | species of plover (all snowy plover 'SNPL' in this case) |
| 2. | population | population at which brood was observed (all Ceuta in this case) |
| 3. | year | year during which brood was observed |
| 4. | site | site at which brood was observed |
| 5. | brood | unique identifier of brood (unique within year and within site). Broods originating from known nests retain the nest identifier found in the *Nests* table, whereas broods hatching from unknown nests have a negative identifier (e.g., -2) |
| 6. | ID | a concatenation of year, site, and nest to make a unique brood ID across sites and years |
| 7. | easting | UTM easting of brood observation |
| 8. | northing | UTM northing of brood observation |
| 9. | utm | UTM zone of brood observation |
| 10. | date | date brood observation was made (stored in the internal 'Date' format of R and represents the number of days since January 1, 1970, the 'Unix epoch'. Converted easily in R using 'as.Date(date, origin = "1970-01-01")') |
| 11. | time | time brood observation was made (24h format) |
| 12. | distance | estimated distance in meters between brood and observer |
| 13. | degree | estimated bearing of brood relative to observer (i.e., the number of degrees in the angle measured in a clockwise direction from the north line to the line joining the observer to the brood) |
| 14. | parents | parents attending brood at time of observation (0 = no parent present; 1 = one parent (not identified whether male or female); 2 = female only (2+ when female certainly identified, whilst male uncertain); 3 = male only (3+, i.e., opposite of 2+); 4 = both present) |
| 15. | male | ring ID of male observed tending brood |
| 16. | female | ring ID of female observed tending brood |
| 17. | chicks | number of chicks observed in brood |
| 18. | chick_codes | color ring combinations of all chicks observed (individuals seperated by a comma). The scheme can be noted as XX.XX\|XX.XX where X indicates a possible position for a color (or metal) ring, the full stop marks the position of 'knee-joint' and the pipe divides the left and right leg. Thus the readout is "left above . left below \| right above . right below". See page 9 of ref. 24 for more details. |
| 19. | photo | indication if a photo of the brood was taken (1) or not (0) |
| 20. | observer | initials of observer making brood observation |
| 21. | comments | miscellaneous comments pertinent to brood's observation |

**Table 4.** Resight data of snowy plovers breeding in Bahía de Ceuta, Mexico, between 2006 and 2016. This dataset contains information on the time and location of a colour-ringed adult, the identity of the individual, and behavioural information recorded during the observation. These data can be used to assess apparent survival with mark-recapture models or investigate space-use through home range analysis or movement ecology models.

| Column | Data name | Description of data |
| --- | --- | --- |
| 1. | species | species of plover (all snowy plover 'SNPL' in this case) |
| 2. | population | population at which resighting was made (all Ceuta in this case) |
| 3. | year | year during which resighting was made |
| 4. | site | site at which resighting was made |
| 5. | easting | UTM easting of observer's location while resighting |
| 6. | northing | UTM northing of observer's location while resighting |
| 7. | utm | UTM zone of observer's location while resighting |
| 8. | date | date resighting was made (stored in the internal 'Date' format of R and represents the number of days since January 1, 1970, the 'Unix epoch'. Converted easily in R using 'as.Date(date, origin = "1970-01-01")') |
| 9. | time | time resighting was made (24h format) |
| 10. | distance | estimated distance in meters between resighted bird and observer |
| 11. | degree | estimated bearing of resighted bird relative to the observer (i.e., the number of degrees in the angle measured in a clockwise direction from the north line to the line joining the observer to the brood) |
| 12. | code | color-ring combination of the resighted individual. The scheme can be noted as XX.XX\|XX.XX where X indicates a possible position for a color (or metal) ring, the full stop marks the position of 'knee-joint' and the pipe divides the left and right leg. Thus the readout is "left above . left below \| right above . right below". See page 9 of ref. 24 for more details. |
| 13. | sex | sex of individual determined in the field based on ornamentation and other clues (e.g., capture history, parental care, etc.), where F = female, M = males, and J = unknown sexed juvenile |
| 14. | census | indication if the resighting was conducted as part of a census count (1) or not (0) |
| 15. | observer | initials of observer making resighting |
| 16. | comments | miscellaneous comments pertinent to the resighting |

**Table 5.** Bird Reference (“BirdRef”) data of snowy plovers breeding in Bahía de Ceuta, Mexico, between 2006 and 2016. This dataset is a relational table of the *Nest data* and *Capture data* summarizing the identity of all members in a family (i.e., colour-rings of both parents and all chicks, if applicable). These data can be used to quantify mating system and assess individual variation in breeding phenology, for example.

| Column | Data name | Description of data |
| --- | --- | --- |
| 1. | species | species of plover (all snowy plover in this case) |
| 2. | population | population at which family was observed (all Ceuta in this case) |
| 3. | year | year during which family was observed |
| 4. | site | site at which family was observed |
| 5. | family | unique identified of family (unique within year and within site). Families found as a nests retain nest ID found in *Nests* table, whereas families found as broods hatching from unknown nests have a negative brood ID (e.g., -2) found in *Broods* table) |
| 6. | ID | a concatenation of year, site, and nest to make a unique family ID across all sites and years |
| 7. | nest_initiation_date | estimated date when the first egg of the nest was laid (i.e., its 'initiation'). The estimate is calculated by subtracting the age in days of the oldest egg (determined by the floatation scores 'float1', 'float2', and 'float3' defined below) and a 5-day laying period for three-egg clutches or a 3-day laying period for two-egg clutches or a 1-day laying period for one-egg clutches (egg-laying intervals are based on ref. 33. Determining initiation dates of clutches found at stage 'F' is imprecise, and thus we estimated the initiation date by subtracting 25 days from the hatch date (i.e., the average length of incubation in this population) and an additional 5, 3, or 1 days for the laying period depending on the clutch size. For nests found at stage 'F' that failed before hatching, the nest initiation date is 'NA') |
| 8. | hatching_date | date nest hatched (stored in the internal 'Date' format of R and represents the number of days since January 1, 1970, the 'Unix epoch'. Converted easily in R using 'as.Date(hatching_date, origin = "1970-01-01")'; "NA" if nest fate was other than "hatch" in *Nests* table) |
| 9. | male | ring ID of male parent observed with nest/brood |
| 10. | female | ring ID of female parent observed with nest/brood |
| 11. | chick1 | ring ID of first chick seen in brood |
| 12. | chick2 | ring ID of second chick seen in brood |
| 13. | chick3 | ring ID of third chick seen in brood |
| 14. | exp | indication if family was part of an experiment |
| 15. | type | indication of type of experiment conducted |
| 16. | manip | date of possible experimental manipulation (stored in the internal 'Date' format of R and represents the number of days since January 1, 1970, the 'Unix epoch'. Converted easily in R using 'as.Date(manip, origin = "1970-01-01")') |

In summary, the CeutaOPEN database contains information on 794 surveyed nests, 3,358 captures of 1,600 marked individuals, 415 monitored broods, and 6,939 resightings of colour-marked individuals. Over the 11-year study period, we spent 927 days collecting these data in the field – amounting to over 20,000 hours of observational effort.

CeutaOPEN is one of only a few open-access databases to provide raw field observations of an individually-marked wild vertebrate species (for other examples, see refs. 30 and 31). We therefore believe our database will provide a valuable model for future field biologists to consult when structuring their data and deciding whether to provide public access.

### Technical Validation

During each field season of the snowy plover project at Bahía de Ceuta, observers receive comprehensive training on our sampling protocol^24^ and general avian field methodology. In all 10 years of data collection included in the CeutaOPEN database presented here, at least one of us was present regularly in the field to oversee fieldwork and assess the quality of observations. Moreover, field assistants usually aided us with fieldwork for academic purposes (e.g., as part of a bachelor, master, or doctoral project), which encouraged personal interest in maximizing the quality of their data collection. Most of the data from the 2014 field season was lost, which is why this year is missing from the database over the 11-year period between 2006 and 2016.

During the data processing and development of the final database, verification and validations were made at several stages: during fieldwork we would regularly check each other’s notes for unusual observations, during the digitization of field data in spreadsheets we would scrutinize outlier measurements, and throughout the assembly of the SQL database we conducted thorough data cleaning (e.g., removing white space from strings, enforcing consistent notation and symbology, etc.). These data quality checks were run annually before merging with the master database.

### Usage Notes

The CeutaOPEN database is available under a Creative Commons Attribution 4.0 International Public License, whereby anyone may freely use and adapt our data, as long as the original source is credited, the original license is linked, and any changes to our data are indicated in subsequent use. Despite our best efforts, the database may include occasional errors or inconsistencies that will be corrected or erased in future versions when we become aware of them. No warranty for accuracy of the data is provided. When using any of the CeutaOPEN materials presented here, please cite this Data Descriptor in addition to the version of the database that was used. Furthermore, for all projects making considerable use of the CeutaOPEN database, we encourage users to reach out to us to offer the opportunity to comment prior to the publication of their work.

We recommend that users employ R to access and wrangle the CeutaOPEN database for their study. To help this process, please refer to the accompanying RMarkdown document (Supplementary File 1) to follow our suggested analytical workflow for utilizing CetuaOPEN with the RSQLite^27^ and dplyr^28^ packages in the R environment.

## Acknowledgements

We thank all of the volunteers in the field for their help monitoring snowy plovers in Ceuta between 2006 and 2016, especially Araceli Argüelles-Tico, Juanita Fonseca-Parra, Cristina Carmona-Isunza, Karla Alavarado-Castro, Raul Said Quintero-Félix, René Beamonte-Barrientos, Oscar Sánchez-Velázquez, David Garcia-Jacome, and Oliva Castañeda-González. We thank Lina Maria Giraldo Deck for statistical advice. We are particularly grateful to Tamás Székely for providing initial funds, protocols, and guidance for the data collection. Xico Vega, Ivan Guardado, Martín A. Serrano-Meneses, and Marcos Búcio-Pacheco helped with permit applications and field logistics. Microsatellite genotyping at NBAF-Sheffield was supported by grants NBAF547, NBAF933, NBAF441. Long-term monitoring would not have been possible without the support of CONACyT (Convocatoria Ciencia Básica SEP-CONACYT 2010 – project number 157570) and the Tracy Aviary’s ‘Dollars to Conservation’ program. For a full list of funding sources please see www.chorlito.org. LJE-P was funded by the German Science Foundation (DFG Eigene Stelle grant: EB 590/1) and the Max Planck Society, MC-L was supported by a doctoral grant from CONACyT (248125/378124) and the Cornell Lab of Ornithology ‘Coastal Solutions Fellowship’, and CK was funded by the Max Planck Society.

## Author contributions

LE-P wrote the Data Descriptor, constructed the database, and coded the accompanying RMarkdown document (Supplementary File 1).

MC-L collected and curated data, and managed field operations.

LLA collected and curated data, and managed field operations.

SGdA collected and curated data, and managed field operations.

WR-A collected and curated data, and managed field operations.

CK initiated the study, collected and curated data, and managed field operations.

All authors helped revise the paper.

## Supplementary Files

Supplementary File 1. An html RMarkdown document providing a detailed commented workflow for utilizing CetuaOPEN with the RSQLite^27^ and dplyr^28^ packages in R. The file can be downloaded and viewed here: https://github.com/leberhartphillips/Ceuta_OPEN/blob/master/R/Ceuta_OPEN.html

