## Supplementary File 1 for "CeutaOPEN: Individual-based field observations of breeding snowy plovers *Charadrius nivosus*": 884825_file02.html

CeutaOPEN


### CeutaOPEN

###### Luke J. Eberhart-Phillips, Medardo Cruz-López, Lydia Lozano-Angulo, Salvador Gómez del Ángel, Wendoly Rojas-Abreu, and Clemens Küpper *Max Planck Institute for Ornithology, Seewiesen, Germany* *Supplementary File 1*

###### 20 December, 2019

In this R Markdown document we provide a simple workflow in R to introduce users to the CeutaOPEN database presented in our paper. In short, the database contains all of our raw field data collected from 1,598 individually marked snowy plovers (*Charadrius nivosus*) monitored between 2006 and 2016 at Bahía de Ceuta – an important breeding site in western Mexico. To access the database and this accompanying R Markdown code, please visit this GitHub repository. An explanation of the datatables in the repository can be found in the **`README.md`** file. Please don’t hesitate to contact Luke Eberhart-Phillips (`luke.eberhart[at]orn.mpg.de`) or Clemens Küpper (`ckuepper[at]orn.mpg.de`) if you have any questions.

The document is best viewed in html on an internet browser as many of the graphics are dynamic.

#### Prerequisites

- To utilize this R Markdown code you need to download the CeutaOPEN GitHub repository containing the **`Ceuta_OPEN.sqlite`** database and other dependent files and folders. Simply follow the link and click on the green *Clone or download* button on the right side of the page and select *Download ZIP*. For a smooth workflow, we recommend creating an RStudio project within the downloaded folder and conducting all further analyses within this directory.
- The following packages are needed to employ this R Markdown document and can be easily installed from CRAN by uncommenting the `install.packages` function below:

```
# install.packages(c("RSQLite", "dplyr", "dbplyr", "knitr", "stringr", "sp", "rgdal", 
#                    "mapview", "RColorBrewer", "webshot", "leaflet", "ggplot2",
#                    "kableExtra", "plotly", "bamlss", "gridExtra"))
# webshot::install_phantomjs()
library(RSQLite)
library(dplyr)
library(dbplyr)
library(knitr)
library(stringr)
library(sp)
library(rgdal)
library(mapview)
library(RColorBrewer)
library(webshot)
library(leaflet)
library(ggplot2)
library(kableExtra)
library(plotly)
```

- The following `as.Date.multicol()` function is useful for transforming date strings within CeutaOPEN tables prior to analysis. Note: date strings in CeutaOPEN are always stored in the internal `Date` format of R, which represents the number of days since January 1, 1970, the ‘Unix epoch’.

```
as.Date.multicol <- function(df){
  if(sum(grepl(paste(c("date", "alive", "manip"), collapse = "|"), names(df))) > 1){
    
    df[, which(grepl(paste(c("date", "alive", "manip"), collapse = "|"), names(df)))] <-
      lapply(df[, which(grepl(paste(c("date", "alive", "manip"), collapse = "|"), names(df)))],
             function(x) as.Date(x, origin = "1970-01-01"))
    
  }
  else{
    
    df[, which(grepl(paste(c("date", "alive", "manip"), collapse = "|"), names(df)))] <- 
      as.Date(df[, which(grepl(paste(c("date", "alive", "manip"), collapse = "|"), 
                               names(df)))], 
              origin = "1970-01-01")
    
  }
  
  return(df)
}
```

#### Data Accessibility

Our database is stored as an SQLite to make it convenient for users to utilize across a variety of platforms. Here we demonstrate how to employ R to access and explore the structure of the database using the ‘RSQLite’ package.

Connect to the **`Ceuta_OPEN`** database

```
Ceuta_OPEN <- RSQLite::dbConnect(SQLite(), dbname="data/Ceuta_OPEN_version_releases/Ceuta_OPEN_v1.sqlite")
```

Our database includes five tables that encompass all of our field observations. Our field methods for obtaining the data presented in these tables is described in the *Methods* section of our paper.

List the tables in the database

```
RSQLite::dbListTables(Ceuta_OPEN)
```

```
## [1] "BirdRef"  "Broods"   "Captures" "Nests"    "Resights"
```

Show the columns in the **`Nests`** table, for example

```
dbListFields(Ceuta_OPEN, "Nests")
```

```
##  [1] "species"                "population"            
##  [3] "year"                   "site"                  
##  [5] "nest"                   "ID"                    
##  [7] "easting"                "northing"              
##  [9] "utm"                    "found_date"            
## [11] "found_time"             "nest_initiation_date"  
## [13] "end_date"               "last_observation_alive"
## [15] "fate"                   "male"                  
## [17] "female"                 "no_chicks"             
## [19] "clutch_size"            "length1"               
## [21] "width1"                 "float1"                
## [23] "length2"                "width2"                
## [25] "float2"                 "length3"               
## [27] "width3"                 "float3"                
## [29] "photo"                  "observer"              
## [31] "comments"
```

#### Making Data Queries

To make database queries, you can use Structured Query Language (SQL) syntax or dplyr syntax. Here we demonstrate how to make a query using both methods, however in subsequent sections of this tutorial we solely employ dplyr as it arguably uses more intuitive syntax for those familiar with the R language.

###### Example 1. *Extract all nests monitored in 2012*

###### *SQL syntax:*

(note that ‘limit 10’ is used here to display only the first 10 observations of the query)

```
kable(dbGetQuery(Ceuta_OPEN, "SELECT * 
                              FROM Nests 
                              WHERE year='2012' 
                              LIMIT 10")) %>%
  kable_styling() %>%
  scroll_box(width = "100%")
```

| species | population | year | site | nest | ID | easting | northing | utm | found\_date | found\_time | nest\_initiation\_date | end\_date | last\_observation\_alive | fate | male | female | no\_chicks | clutch\_size | length1 | width1 | float1 | length2 | width2 | float2 | length3 | width3 | float3 | photo | observer | comments |
| --- | --- | --- | --- | --- | --- | --- | --- | --- | --- | --- | --- | --- | --- | --- | --- | --- | --- | --- | --- | --- | --- | --- | --- | --- | --- | --- | --- | --- | --- | --- |
| SNPL | Ceuta | 2012 | G | 301 | 2012\_G\_301 | 299329 | 2647323 | NA | 15487 | NA | 15474 | 15495 | NA | Flooded | NA | NA | 0 | 3 | 30.7 | 22.7 | D | 30.2 | 23.1 | C | 31.0 | 23.1 | C | NA | MC | NA |
| SNPL | Ceuta | 2012 | D | 4 | 2012\_D\_4 | 299808 | 2646480 | NA | 15475 | NA | 15462 | 15493 | NA | Hatch | NA | NA | 3 | 3 | 32.0 | 23.5 | C | 31.5 | 23.1 | D | 31.5 | 23.5 | D | NA | CK | NA |
| SNPL | Ceuta | 2012 | G | 201 | 2012\_G\_201 | 299299 | 2647323 | NA | 15480 | NA | 15467 | 15495 | NA | Flooded | NA | NA | 0 | 3 | 32.0 | 22.7 | B | 31.5 | 23.0 | C | 31.2 | 22.2 | D | NA | AS | NA |
| SNPL | Ceuta | 2012 | C | 301 | 2012\_C\_301 | 300531 | 2646195 | NA | 15503 | NA | 15490 | NA | NA | Unknown | NA | NA | NA | 3 | 32.4 | 22.8 | C | 30.5 | 23.0 | D | 29.9 | 22.8 | C | NA | MC | NA |
| SNPL | Ceuta | 2012 | G | 204 | 2012\_G\_204 | 299273 | 2647213 | NA | 15482 | NA | 15472 | 15495 | NA | Flooded | NA | NA | 0 | 3 | 31.8 | 22.8 | C | 32.5 | 22.8 | C | 32.9 | 22.5 | B | NA | AS | NA |
| SNPL | Ceuta | 2012 | G | 305 | 2012\_G\_305 | 299352 | 2647286 | NA | 15494 | NA | 15484 | 15495 | NA | Flooded | NA | NA | 0 | 3 | 29.4 | 22.3 | C | 31.2 | 22.8 | B | 30.9 | 22.6 | B | NA | MC | Parents not captured |
| SNPL | Ceuta | 2012 | D | 102 | 2012\_D\_102 | 300142 | 2646862 | NA | 15491 | NA | 15484 | 15515 | NA | Predated | NA | NA | 0 | 3 | 30.6 | 22.9 | B | 30.7 | 22.8 | B | 30.6 | 22.7 | AB | NA | WR | predated by racoon |
| SNPL | Ceuta | 2012 | G | 1 | 2012\_G\_1 | 299242 | 2647369 | NA | 15467 | NA | 15460 | 15493 | NA | Hatch | NA | NA | 1 | 3 | 32.2 | 23.1 | B | 31.5 | 22.8 | A | 31.9 | 23.0 | AB | NA | CK | NA |
| SNPL | Ceuta | 2012 | G | 202 | 2012\_G\_202 | 299325 | 2647530 | NA | 15480 | NA | 15470 | 15495 | NA | Flooded | NA | NA | 0 | 3 | 32.1 | 22.5 | B | 31.7 | 22.7 | B | 32.9 | 22.1 | C | NA | AS | NA |
| SNPL | Ceuta | 2012 | D | 1 | 2012\_D\_1 | 299823 | 2646570 | NA | 15458 | NA | 15457 | 15489 | NA | Hatch | NA | NA | 3 | 3 | 31.7 | 22.5 | B | 31.2 | 22.6 | C | 30.3 | 21.8 | AB | NA | CK | found with one egg only, all eggs floated 507 |

###### *dplyr syntax:*

```
dbReadTable(Ceuta_OPEN, "Nests") %>%
  filter(year == 2012) %>%
  head(10) %>%
  kable() %>%
  kable_styling() %>%
  scroll_box(width = "100%")
```

| species | population | year | site | nest | ID | easting | northing | utm | found\_date | found\_time | nest\_initiation\_date | end\_date | last\_observation\_alive | fate | male | female | no\_chicks | clutch\_size | length1 | width1 | float1 | length2 | width2 | float2 | length3 | width3 | float3 | photo | observer | comments |
| --- | --- | --- | --- | --- | --- | --- | --- | --- | --- | --- | --- | --- | --- | --- | --- | --- | --- | --- | --- | --- | --- | --- | --- | --- | --- | --- | --- | --- | --- | --- |
| SNPL | Ceuta | 2012 | G | 301 | 2012\_G\_301 | 299329 | 2647323 | NA | 15487 | NA | 15474 | 15495 | NA | Flooded | NA | NA | 0 | 3 | 30.7 | 22.7 | D | 30.2 | 23.1 | C | 31.0 | 23.1 | C | NA | MC | NA |
| SNPL | Ceuta | 2012 | D | 4 | 2012\_D\_4 | 299808 | 2646480 | NA | 15475 | NA | 15462 | 15493 | NA | Hatch | NA | NA | 3 | 3 | 32.0 | 23.5 | C | 31.5 | 23.1 | D | 31.5 | 23.5 | D | NA | CK | NA |
| SNPL | Ceuta | 2012 | G | 201 | 2012\_G\_201 | 299299 | 2647323 | NA | 15480 | NA | 15467 | 15495 | NA | Flooded | NA | NA | 0 | 3 | 32.0 | 22.7 | B | 31.5 | 23.0 | C | 31.2 | 22.2 | D | NA | AS | NA |
| SNPL | Ceuta | 2012 | C | 301 | 2012\_C\_301 | 300531 | 2646195 | NA | 15503 | NA | 15490 | NA | NA | Unknown | NA | NA | NA | 3 | 32.4 | 22.8 | C | 30.5 | 23.0 | D | 29.9 | 22.8 | C | NA | MC | NA |
| SNPL | Ceuta | 2012 | G | 204 | 2012\_G\_204 | 299273 | 2647213 | NA | 15482 | NA | 15472 | 15495 | NA | Flooded | NA | NA | 0 | 3 | 31.8 | 22.8 | C | 32.5 | 22.8 | C | 32.9 | 22.5 | B | NA | AS | NA |
| SNPL | Ceuta | 2012 | G | 305 | 2012\_G\_305 | 299352 | 2647286 | NA | 15494 | NA | 15484 | 15495 | NA | Flooded | NA | NA | 0 | 3 | 29.4 | 22.3 | C | 31.2 | 22.8 | B | 30.9 | 22.6 | B | NA | MC | Parents not captured |
| SNPL | Ceuta | 2012 | D | 102 | 2012\_D\_102 | 300142 | 2646862 | NA | 15491 | NA | 15484 | 15515 | NA | Predated | NA | NA | 0 | 3 | 30.6 | 22.9 | B | 30.7 | 22.8 | B | 30.6 | 22.7 | AB | NA | WR | predated by racoon |
| SNPL | Ceuta | 2012 | G | 1 | 2012\_G\_1 | 299242 | 2647369 | NA | 15467 | NA | 15460 | 15493 | NA | Hatch | NA | NA | 1 | 3 | 32.2 | 23.1 | B | 31.5 | 22.8 | A | 31.9 | 23.0 | AB | NA | CK | NA |
| SNPL | Ceuta | 2012 | G | 202 | 2012\_G\_202 | 299325 | 2647530 | NA | 15480 | NA | 15470 | 15495 | NA | Flooded | NA | NA | 0 | 3 | 32.1 | 22.5 | B | 31.7 | 22.7 | B | 32.9 | 22.1 | C | NA | AS | NA |
| SNPL | Ceuta | 2012 | D | 1 | 2012\_D\_1 | 299823 | 2646570 | NA | 15458 | NA | 15457 | 15489 | NA | Hatch | NA | NA | 3 | 3 | 31.7 | 22.5 | B | 31.2 | 22.6 | C | 30.3 | 21.8 | AB | NA | CK | found with one egg only, all eggs floated 507 |

###### Example 2. *List the years and weights from all captures of the bird with the colour-ring combination “BX.MX|RX.RX”*

###### *SQL syntax:*

```
dbGetQuery(Ceuta_OPEN, "SELECT code, year, weight 
                        FROM Captures 
                        WHERE code = 'BX.MX|RX.RX'") %>%
  kable(col.names = c("Colour combination",
                      "Year",
                      "Body mass (g)")) %>%
  kable_styling() %>%
  scroll_box(width = "100%")
```

| Colour combination | Year | Body mass (g) |
| --- | --- | --- |
| BX.MX\|RX.RX | 2008 | 35.80 |
| BX.MX\|RX.RX | 2008 | 36.60 |
| BX.MX\|RX.RX | 2012 | 34.00 |
| BX.MX\|RX.RX | 2011 | 35.40 |
| BX.MX\|RX.RX | 2009 | 37.30 |
| BX.MX\|RX.RX | 2007 | 34.40 |
| BX.MX\|RX.RX | 2011 | 34.50 |

###### *dplyr syntax:*

```
dbReadTable(Ceuta_OPEN, "Captures") %>%
  filter(code == "BX.MX|RX.RX") %>%
  select(code, year, weight) %>%
  collect() %>%
  kable(col.names = c("Colour combination",
                      "Year",
                      "Body mass (g)")) %>%
  kable_styling() %>%
  scroll_box(width = "100%")
```

| Colour combination | Year | Body mass (g) |
| --- | --- | --- |
| BX.MX\|RX.RX | 2008 | 35.80 |
| BX.MX\|RX.RX | 2008 | 36.60 |
| BX.MX\|RX.RX | 2012 | 34.00 |
| BX.MX\|RX.RX | 2011 | 35.40 |
| BX.MX\|RX.RX | 2009 | 37.30 |
| BX.MX\|RX.RX | 2007 | 34.40 |
| BX.MX\|RX.RX | 2011 | 34.50 |

###### Example 3. *Find the top 6 males that have had the most hatched nests over the study period*

###### *SQL syntax:*

```
dbGetQuery(Ceuta_OPEN, "SELECT ring, count(*) as count
                        FROM Nests a
                        JOIN Captures b
                        ON a.ID = b.ID
                        AND a.fate = 'Hatch'
                        AND b.sex = 'M'
                        GROUP BY b.ring
                        ORDER BY count(*) DESC
                        LIMIT 6") %>%
  kable(col.names = c("Ring number",
                      "Number of hatched nests")) %>%
  kable_styling() %>%
  scroll_box(width = "100%")
```

| Ring number | Number of hatched nests |
| --- | --- |
| CA1272 | 8 |
| CA168 | 8 |
| CA078 | 7 |
| CA1203 | 7 |
| CA1213 | 7 |
| CA1310 | 7 |

###### *dplyr syntax:*

```
left_join(x = dbReadTable(Ceuta_OPEN, "Nests"), 
          y = dbReadTable(Ceuta_OPEN, "Captures"), 
          by = "ID") %>%
  filter(sex == "M" & fate == "Hatch") %>%
  group_by(ring) %>%
  tally(sort = TRUE) %>%
  top_n(6) %>%
  collect() %>%
  kable(col.names = c("Ring number",
                      "Number of hatched nests")) %>%
  kable_styling() %>%
  scroll_box(width = "100%")
```

| Ring number | Number of hatched nests |
| --- | --- |
| CA1272 | 8 |
| CA168 | 8 |
| CA078 | 7 |
| CA1203 | 7 |
| CA1213 | 7 |
| CA1310 | 7 |

#### Database Summary

Here we summarise some of the key components from each table in the database.

##### Nests datatable

###### *Number of nests monitored*

```
dbReadTable(Ceuta_OPEN, "Nests") %>%
  summarise(n_nests = n_distinct(ID)) %>%
  kable(col.names = c("Number of nests monitored")) %>%
  kable_styling() %>%
  scroll_box(width = "100%")
```

Number of nests monitored

794

###### *Map of all nests in the study area, coloured by year*

```
# Extract the Nests datatable
Nests <- dbReadTable(Ceuta_OPEN, "Nests")

# Classify the 'easting' and 'northing' columns as numeric
Nests[,c("easting", "northing")] <- 
  lapply(Nests[,c("easting", "northing")], as.numeric)

# Remove nests without spatial coordinates
Nests <- filter(Nests, !is.na(easting))

# Set the UTM 1983 zone 13 North (format nest data is in)
UTM13n <- CRS("+proj=utm +zone=13 +ellps=WGS84 +datum=WGS84 +units=m +no_defs")

# Set the World Geographic System 1984 (lat/long, i.e., format for mapping)
WGS84 <- CRS("+proj=longlat +ellps=WGS84 +datum=WGS84") 

# Create a spatialpointsdataframe while specifying the coordinates
Nests_spat_UTM <-  
  SpatialPointsDataFrame(coords = Nests[,c("easting", "northing")],
                         data = Nests, 
                         proj4string = UTM13n)

# transform the UTM coordinates to lat/long for mapping
Nests_spat_WGS84 <- spTransform(Nests_spat_UTM, WGS84)

# specify mapview options for viewing
mapviewOptions(basemaps = c("Esri.WorldImagery"),
               layers.control.pos = "topright",
               legend.pos = "bottomleft")

# open mapview leaflet of the spatial extent of the Nests data (Note: this is best viewed in an HTML version of this R Markdown document)
nest_map <- 
  mapview(Nests_spat_WGS84, zcol = "year", 
          col.regions = colorRampPalette(brewer.pal(9, "Blues")),
          layer.name = "Snowy Plover Nests",
          layers.control.pos = "topright")

mean_coords <- c(mean(coordinates(Nests_spat_WGS84)[, 1]-.02),
               mean(coordinates(Nests_spat_WGS84)[, 2]))

nest_map@map %>% setView(mean_coords[1], mean_coords[2], zoom = 14)
```

###### *Annual variation in nesting activity and fates*

```
dbReadTable(Ceuta_OPEN, "Nests") %>% 
  group_by(year, fate) %>%
  dplyr::summarise(n_nests = dplyr::n()) %>% 
  mutate(fate = factor(fate, levels = c("Hatch", "Predated", "Flooded", 
                                        "Abandoned", "Unhatched", "Other", 
                                        "Unknown"))) %>% 
  ggplot(aes(y = n_nests, x = year, fill = fate, color = fate)) +
  geom_bar(stat = "identity") +
  scale_color_brewer(palette = "Set2",
                     name = "Nest fate") +
  scale_fill_brewer(palette = "Set2",
                    name = "Nest fate") +
  ylab("Nests monitored")
```

###### *Check the distribution of egg morphometrics*

```
egg_plot(dbReadTable(Ceuta_OPEN, "Nests"), species_name = "SNPL")
```

##### Captures datatable

###### *Number of captures for males, females, and chicks*

```
dbReadTable(Ceuta_OPEN, "Captures") %>%
  group_by(sex, age) %>%
  dplyr::summarise(n_captures = dplyr::n()) %>%
  kable(col.names = c("Sex", "Age group", "Number of captures")) %>%
  kable_styling() %>%
  scroll_box(width = "100%")
```

| Sex | Age group | Number of captures |
| --- | --- | --- |
| F | A | 697 |
| F | J | 633 |
| M | A | 638 |
| M | J | 548 |
| U | J | 253 |

###### *Number of unique individuals in marked population*

```
dbReadTable(Ceuta_OPEN, "Captures") %>%
  dplyr::summarise(n_marked_individuals = n_distinct(ring)) %>%
  kable(col.names = c("Number of individuals ringed")) %>%
  kable_styling() %>%
  scroll_box(width = "100%")
```

Number of individuals ringed

1600

###### *Number of unique individual males, females, and juveniles ringed*

```
dbReadTable(Ceuta_OPEN, "Captures") %>%
  filter(year != 2017 & year != 2018 & year != 2019) %>%
  group_by(sex, age) %>%
  summarise(n_individuals = n_distinct(ring)) %>%
  kable(col.names = c("Sex", "Age group", "Number of captures")) %>%
  kable_styling() %>%
  scroll_box(width = "100%")
```

| Sex | Age group | Number of captures |
| --- | --- | --- |
| F | A | 334 |
| F | J | 441 |
| M | A | 310 |
| M | J | 402 |
| U | J | 195 |

###### *Annual variation in captures*

```
annual_captures <- 
  dbReadTable(Ceuta_OPEN, "Captures") %>%
  group_by(year, sex) %>%
  dplyr::summarise(n_capts = dplyr::n_distinct(ring),
            total_capts = dplyr::n()) %>%
  ungroup() %>%
  mutate(n_capts = as.numeric(n_capts),
         year = as.numeric(year),
         sex = factor(sex, levels = c("F", "M", "U"))) %>%
  collect()

ggplot2::ggplot(data = annual_captures, aes(y = n_capts, x = year, color = sex, fill = sex)) +
  geom_bar(stat = "identity", alpha = 0.7) +
  scale_color_manual(values = plot_palette_sex,
                     name = "Age and sex",
                     breaks = c("F", "M", "U"),
                     labels = c("Adult females", "Adult males", "Juveniles")) +
  scale_fill_manual(values = plot_palette_sex,
                    name = "Age and sex",
                    breaks = c("F", "M", "U"),
                    labels = c("Adult females", "Adult males", "Juveniles")) +
  ylab("Individuals captured") +
  scale_x_continuous(labels = as.character(annual_captures$year), 
                     breaks = annual_captures$year)
```

###### *Check the distribution of body measurments*

```
tarsus_plot(df = dbReadTable(Ceuta_OPEN, "Captures"), species_name = "SNPL")
```

```
wing_plot(df = dbReadTable(Ceuta_OPEN, "Captures"), species_name = "SNPL")
```

```
weight_plot(df = dbReadTable(Ceuta_OPEN, "Captures"), species_name = "SNPL")
```

```
bill_plot(df = dbReadTable(Ceuta_OPEN, "Captures"), species_name = "SNPL")
```

##### Broods datatable

###### *Number of broods monitored*

```
dbReadTable(Ceuta_OPEN, "Broods") %>%
  dplyr::summarise(n_broods = n_distinct(ID)) %>%
  kable(col.names = c("Number of broods monitored")) %>%
  kable_styling() %>%
  scroll_box(width = "100%")
```

Number of broods monitored

415

###### *Annual variation in broods*

```
annual_broods <- 
  dbReadTable(Ceuta_OPEN, "Broods") %>% 
  as.Date.multicol() %>% 
  left_join(as.Date.multicol(dbReadTable(Ceuta_OPEN, "BirdRef")), by = "ID") %>% 
  filter(!is.na(date)) %>% 
  mutate(brood_age = ifelse(date-hatch_date < 0, 0, date-hatch_date)) %>% 
  group_by(year.x, ID) %>%
  dplyr::summarise(max_age = max(brood_age)) %>%
  filter(!is.na(max_age)) %>%
  mutate(week_bin = ifelse(max_age < 8, "1 week",
         ifelse(max_age > 7 & max_age < 15, "2 weeks",
                ifelse(max_age > 14 & max_age < 22, "3 weeks",
                       ifelse(max_age > 21 & max_age < 29, "4 weeks",
                              ifelse(max_age > 28 , "5+ weeks", "XXX")))))) %>%
  ungroup() %>% 
  mutate(year.x = as.factor(year.x),
         week_bin = as.factor(week_bin)) %>%
  dplyr::group_by(year.x, week_bin) %>%
  dplyr::summarise(n_brods = n_distinct(ID)) %>%
  ungroup() %>% 
  mutate(n_brods = as.numeric(n_brods),
         year.x = as.numeric(as.character(year.x)))

ggplot2::ggplot(data = annual_broods, aes(y = n_brods, x = year.x, color = week_bin, fill = week_bin)) +
  geom_bar(stat = "identity", alpha = 0.7) +
  scale_fill_manual(values = plot_palette_brood,
                    name = "Max age\nobserved") +
  scale_color_manual(values = plot_palette_brood,
                     name = "Max age\nobserved") +
  ylab("Broods monitored") +
  scale_x_continuous(labels = as.character(annual_broods$year.x), breaks = annual_broods$year.x)
```

###### *average number of resightings per brood*

```
dbReadTable(Ceuta_OPEN, "Broods") %>%
  group_by(ID) %>%
  dplyr::summarise(n_obs_per_brood = dplyr::n()) %>%
  dplyr::summarise(mean = mean(n_obs_per_brood)) %>%
  kable(col.names = c("Average number of resightings per brood")) %>%
  kable_styling() %>%
  scroll_box(width = "100%")
```

Average number of resightings per brood

8.250602

###### *Average interval in days between brood resightings*

```
dbReadTable(Ceuta_OPEN, "Broods") %>% 
  as.Date.multicol() %>% 
  dplyr::group_by(year, ID) %>%
  dplyr::summarise(n_obs_per_brood = dplyr::n(),
                   min_date = min(date),
                   max_date = max(date)) %>%
  dplyr::mutate(diff_days = max_date - min_date) %>%
  dplyr::mutate(avg_days = diff_days/n_obs_per_brood) %>%
  dplyr::ungroup() %>%
  dplyr::summarise(avg_days_per_brood = mean(avg_days, na.rm = TRUE)) %>%
  kable(col.names = c("Average interval in days between brood resightings")) %>%
  kable_styling() %>%
  scroll_box(width = "100%")
```

Average interval in days between brood resightings

1.89469 days

##### Resights datatable

###### *Number of resightings*

```
dbReadTable(Ceuta_OPEN, "Resights") %>%
  dplyr::summarise(n_resightings = dplyr::n()) %>%
  kable(col.names = c("Total number of resightings")) %>%
  kable_styling() %>%
  scroll_box(width = "100%")
```

Total number of resightings

6939

###### *Average number of resightings per individual over their observed lifetime in the population*

```
dbReadTable(Ceuta_OPEN, "Resights") %>%
  group_by(code) %>% 
  dplyr::summarise(n_resightings_per_bird = dplyr::n()) %>%
  dplyr::summarise(mean = mean(n_resightings_per_bird)) %>% 
  kable(col.names = c("Average number of resightings per individual")) %>%
  kable_styling() %>%
  scroll_box(width = "100%")
```

Average number of resightings per individual

11.17391

##### BirdRef datatable

###### *Number of unique families monitored*

```
dbReadTable(Ceuta_OPEN, "BirdRef") %>%
  dplyr::summarise(n_families = n_distinct(ID)) %>%
  kable(col.names = c("Number of families monitored")) %>%
  kable_styling() %>%
  scroll_box(width = "100%")
```

Number of families monitored

827

###### *Number of nests discovered after hatch*

(Note: these have negative nest IDs to indicate that they were initially found as broods)

```
dbReadTable(Ceuta_OPEN, "BirdRef") %>%
  filter(str_detect(string = nest, pattern = "-")) %>%
  group_by(year) %>% 
  summarise(n_neg_fams = n_distinct(ID))
```

```
## # A tibble: 8 x 2
##   year  n_neg_fams
##   <chr>      <int>
## 1 2006           4
## 2 2007           8
## 3 2008           3
## 4 2009           2
## 5 2010          12
## 6 2013           1
## 7 2015           4
## 8 2016          10
```

###### *Proportion of families with both, either, or no parents identified*

```
dbReadTable(Ceuta_OPEN, "BirdRef") %>%
  mutate(completeness = ifelse(!is.na(male) & !is.na(female), "both", 
                               ifelse(!is.na(male) & is.na(female), "male",
                                      ifelse(is.na(male) & !is.na(female), "female",
                                             ifelse(is.na(male) & is.na(female), "neither",
                                                    "XXXX"))))) %>% 
  group_by(completeness) %>% 
  summarise(tally = dplyr::n()) %>%
  kable(col.names = c("Type", "Frequency")) %>%
  kable_styling() %>%
  scroll_box(width = "100%")
```

| Type | Frequency |
| --- | --- |
| both | 560 |
| female | 93 |
| male | 46 |
| neither | 128 |

#### Example Analytical Workflow

##### Sex-specific Ontogeny

In this section we provide an example workflow of how to use the CeutaOPEN database in R to assess sex differences in chick developement from hatching until fledging. In the field, we attempt to recapture uniquely marked chicks repeatedly until they fledge to assess their condition by measuring their tarsi (i.e., the lower leg segment) and body mass. These individuals are also molecularly sexed from a small blood sample that is collected during their first capture - allowing a confident assessment of sex-specific growth rates.

###### 1) Load R packages

```
# RSQLite is needed to bring the SQL database into the R environment
library(RSQLite)

# dplyr is useful for wrangling data prior to analysis
library(dplyr)

# ggplot is useful for data visualization
library(ggplot2)

# bamlss is used for "Bayesian Additive Models for Location, Scale, and Shape"
# and is needed here to determine the sex-specific growth trends while controlling
# for repeated measures within individuals and random annual variation
library(bamlss)

# gridExtra allows us to create a beautiful custom layout for plotting the results
library(gridExtra)

# lubridate enables the use of simple functions related to time strings
library(lubridate)

# stringr offers useful functions to manipulate character srings
library(stringr)
```

###### 2) Custom Functions

Convert all columns with date into the `%Y-%m-%d` format. Note the the current format of CeutaOPEN tables is the internal `Date` format of R and represents the number of days since January 1, 1970, the ‘Unix epoch’ (i.e., “the 0-second of one of Humankind’s first computer operating systems”)

```
# date conversion function (columns with `date` or `alive` in their header are 
# converted to the `%Y-%m-%d` format)
as.Date.multicol <- function(df){
  if(sum(grepl(paste(c("date", "alive", "manip"), collapse = "|"), names(df))) > 1){
    
    df[, which(grepl(paste(c("date", "alive", "manip"), collapse = "|"), names(df)))] <-
      lapply(df[, which(grepl(paste(c("date", "alive", "manip"), collapse = "|"), names(df)))],
             function(x) as.Date(x, origin = "1970-01-01"))
    
  }
  else{
    
    df[, which(grepl(paste(c("date", "alive", "manip"), collapse = "|"), names(df)))] <- 
      as.Date(df[, which(grepl(paste(c("date", "alive", "manip"), collapse = "|"), 
                               names(df)))], 
              origin = "1970-01-01")
    
  }
  
  return(df)
}
```

###### 3) Load CeutaOPEN into R

Load the CeutaOPEN database into R using the RSQLite package

```
Ceuta_OPEN <- RSQLite::dbConnect(SQLite(), dbname = "data/Ceuta_OPEN_version_releases/Ceuta_OPEN_v1.sqlite")
```

###### 4) Data Wrangling

In this case we need to wrangle the `Captures` table and extract all cases in which a chick has been captured more than once, calculate the average tarsus length based on the left and right tarsal measurement in each capture, and convert the date column into the appropriate format discussed in step 3 above.

```
multi_cap_chicks <-
  # read the Captures table
  dbReadTable(Ceuta_OPEN, "Captures") %>% 
  # average the left and right tarsus length measurements in to one value
  mutate(tarsus = rowMeans(cbind(as.numeric(left_tarsus), 
                                 as.numeric(right_tarsus)), na.rm = TRUE)) %>% 
  # subset to juvenile captures
  filter(age == "J",
  # remove observations that are missing information for weight and tarsus
         !is.na(weight) & !is.na(tarsus),
  # subset to individuals for which we have the determined the sex molecularly
         sex %in% c("M", "F")) %>% 
  # group by bird identity
  group_by(ring) %>% 
  # subset to juveniles with more than 1 capture
  filter(dplyr::n() > 1) %>% 
  # specifiy the biological nest ID as the ID of the earliest capture (i.e.,
  # brood mixing can occur and create multiple nest IDs for a chick in the
  # capture data)
  mutate(bio_ID = ID[which.min(date)]) %>% 
  # filter(bio_ID != ID) %>% 
  # convert to dataframe
  data.frame() %>% 
  # convert date columns to the %Y-%m-%d format
  as.Date.multicol() %>% 
  # select the relevent columns
  select(ring, year, bio_ID, age, sex, date, time, weight, tarsus)
```

Wrangle the `Nests` table to extract the hatch dates of each of the chicks isolated in the previous step.

```
hatch_dates <- 
  # read the Nests table
  dbReadTable(Ceuta_OPEN, "Nests") %>% 
  # extract only the nests that contain chicks in the previous capture subset
  filter(ID %in% multi_cap_chicks$bio_ID) %>% 
  # subset the nests that have hatch date information
  filter(fate == "Hatch") %>% 
  # define as a dataframe
  data.frame() %>% 
  # classify date columns in the appropriate format
  as.Date.multicol() %>%
  # subset the result as simply the nest ID and their resepective hatch dates
  select(ID, end_date) %>% 
  # rename the columns
  rename(bio_ID = ID,
         hatch_date = end_date)
```

Join the chick capture history to the hatch dates and prep variables for modelling

```
chicks_and_hatch_dates <- 
  # join the hatch dates to the chick captures
  left_join(x = multi_cap_chicks, y = hatch_dates, by = "bio_ID") %>%
  # classify variables as factor or numeric and calculate age at capture
  ungroup() %>% 
  mutate(age = as.numeric(date - hatch_date),
         year = as.factor(year),
         weight = as.numeric(weight),
         tarsus = as.numeric(tarsus),
         ring = as.factor(ring),
         sex = as.factor(sex)) %>% 
  # specify ages less than 0 as 0 (i.e., hatch dates represent the average hatch
  # date of a brood and thus the earliest chick to hatch in a nest could be up
  # 2 days earlier than the nest's hatch date)
  mutate(age = ifelse(age < 0, 0, age)) %>% 
  # scale the numeric variables in preparation for modelling
  mutate(age.z = scale(age),
         weight.z = scale(weight),
         tarsus.z = scale(tarsus)) %>% 
  # remove individuals that don't have an age value (i.e., hatch date was NA)
  filter(!is.na(age))
```

Create sex-specific dataframes

```
female_chicks <- 
  chicks_and_hatch_dates %>%
  # subset to females
  filter(sex == "F") 

male_chicks <- 
  chicks_and_hatch_dates %>% 
  # subset to males
  filter(sex == "M")
```

Wrangle the `Captures` table to extract the adult weights of the chicks used in the analysis above

```
adult_measurements <- 
  dbReadTable(Ceuta_OPEN, "Captures") %>%
  # subset to adults that have the same ring as the chicks indentified earlier
  filter(ring %in% chicks_and_hatch_dates$ring) %>% 
  filter(age == "A") %>% 
  # calculate the average tarsus length across multiple adult captures
  mutate(left_tarsus = as.numeric(left_tarsus),
         right_tarsus = as.numeric(right_tarsus),
         weight = as.numeric(weight)) %>% 
  group_by(ring, sex) %>% 
  summarise(weight = mean(weight, na.rm = TRUE),
            left_tarsus = mean(left_tarsus, na.rm = TRUE),
            right_tarsus = mean(right_tarsus, na.rm = TRUE)) %>% 
  mutate(tarsus = rowMeans(cbind(left_tarsus, right_tarsus), na.rm = TRUE),
         age = "A") %>% 
  select(ring, weight, tarsus, age, sex)
```

###### 5) Modelling

Run models to estimate trend lines of tarsus growth and body mass change over age using a Bayesian generalized additive model with year and individual as random effects.

```
weight_mod_F <- bamlss(weight.z ~ s(age.z) + s(ring, bs = "re") + 
                         s(year, bs = "re") + s(ring, age.z, bs = "re"), 
                       data = female_chicks)
tarsus_mod_F <- bamlss(tarsus.z ~ s(age.z) + s(ring, bs = "re") + 
                         s(year, bs = "re") + s(ring, age.z, bs = "re"), 
                       data = female_chicks)
weight_mod_M <- bamlss(weight.z ~ s(age.z) + s(ring, bs = "re") + 
                         s(year, bs = "re") + s(ring, age.z, bs = "re"), 
                       data = male_chicks)
tarsus_mod_M <- bamlss(tarsus.z ~ s(age.z) + s(ring, bs = "re") + 
                         s(year, bs = "re") + s(ring, age.z, bs = "re"), 
                       data = male_chicks)
```

Extract the mean and sd statistics and re-scale the model coefficients - including the upper and lower 95% credible intervals

```
weight.z_center_F <- attributes(female_chicks$weight.z)$`scaled:center`
weight.z_scale_F <- attributes(female_chicks$weight.z)$`scaled:scale`
tarsus.z_center_F <- attributes(female_chicks$tarsus.z)$`scaled:center`
tarsus.z_scale_F <- attributes(female_chicks$tarsus.z)$`scaled:scale`

female_chicks_output <- 
  female_chicks %>% 
  mutate(lwrm_w = fitted(weight_mod_F, term = c("s(age.z)"))$mu[, 1],
         fitm_w = fitted(weight_mod_F, term = c("s(age.z)"))$mu[, 2],
         uprm_w = fitted(weight_mod_F, term = c("s(age.z)"))$mu[, 3],
         lwrm_t = fitted(tarsus_mod_F, term = c("s(age.z)"))$mu[, 1],
         fitm_t = fitted(tarsus_mod_F, term = c("s(age.z)"))$mu[, 2],
         uprm_t = fitted(tarsus_mod_F, term = c("s(age.z)"))$mu[, 3]) %>% 
  mutate(fitm33_w = fitm_w * weight.z_scale_F + weight.z_center_F,
         lwrm33_w = lwrm_w * weight.z_scale_F + weight.z_center_F,
         uprm33_w = uprm_w * weight.z_scale_F + weight.z_center_F,
         fitm33_t = fitm_t * tarsus.z_scale_F + tarsus.z_center_F,
         lwrm33_t = lwrm_t * tarsus.z_scale_F + tarsus.z_center_F,
         uprm33_t = uprm_t * tarsus.z_scale_F + tarsus.z_center_F) %>% 
  arrange(age)

weight.z_center_M <- attributes(male_chicks$weight.z)$`scaled:center`
weight.z_scale_M <- attributes(male_chicks$weight.z)$`scaled:scale`
tarsus.z_center_M <- attributes(male_chicks$tarsus.z)$`scaled:center`
tarsus.z_scale_M <- attributes(male_chicks$tarsus.z)$`scaled:scale`

male_chicks_output <- 
  male_chicks %>% 
  mutate(lwrm_w = fitted(weight_mod_M, term = c("s(age.z)"))$mu[, 1],
         fitm_w = fitted(weight_mod_M, term = c("s(age.z)"))$mu[, 2],
         uprm_w = fitted(weight_mod_M, term = c("s(age.z)"))$mu[, 3],
         lwrm_t = fitted(tarsus_mod_M, term = c("s(age.z)"))$mu[, 1],
         fitm_t = fitted(tarsus_mod_M, term = c("s(age.z)"))$mu[, 2],
         uprm_t = fitted(tarsus_mod_M, term = c("s(age.z)"))$mu[, 3]) %>% 
  mutate(fitm33_w = fitm_w * weight.z_scale_M + weight.z_center_M,
         lwrm33_w = lwrm_w * weight.z_scale_M + weight.z_center_M,
         uprm33_w = uprm_w * weight.z_scale_M + weight.z_center_M,
         fitm33_t = fitm_t * tarsus.z_scale_M + tarsus.z_center_M,
         lwrm33_t = lwrm_t * tarsus.z_scale_M + tarsus.z_center_M,
         uprm33_t = uprm_t * tarsus.z_scale_M + tarsus.z_center_M) %>% 
  arrange(age)

chick_growth_results <- 
  bind_rows(female_chicks_output, male_chicks_output)
```

###### 7) Plotting

Plot the sex-specific ontenigenic change in tarsus length and body mass over age while comparing the end result to the adult distributions

```
# define the color palatte to use for visualizing males and females
sex_palette <- brewer.pal(7, "Dark2")[c(2,1)]

# draw the chick tarsus plot
chick_tarsus_plot <- 
  ggplot() +
  geom_line(data = chick_growth_results, aes(y = fitm33_t, x = age, 
                                      color = sex),
            size = 1) +
  geom_ribbon(data = chick_growth_results, aes(ymin = uprm33_t, ymax = lwrm33_t, x = age, 
                                        fill = sex), alpha = 0.3) +
  geom_point(data = chick_growth_results, aes(y = tarsus, x = age, 
                                       fill = sex, 
                                       color = sex),
             alpha = 0.3, size = 2) +
  ylab("Tarsus length ± 95% CI (mm)") +
  xlab("Days since hatching") +
  scale_color_manual(values = sex_palette,
                     name = "Sex",
                     labels = c("Female", "Male")) +
  scale_fill_manual(values = sex_palette,
                    name = "Sex",
                    labels = c("Female", "Male")) +
  theme(legend.position = c(0.2, 0.8),
        axis.title.x = element_blank(),
        axis.text.x  = element_blank()) +
  scale_y_continuous(limits = c(15, 30))

# draw the adult tarsus plot
adult_tarsus_plot <- 
  ggplot(adult_measurements, aes(tarsus)) + 
  geom_density(aes(fill = sex), alpha = 0.3, color = "grey40") + 
  scale_fill_manual(values = sex_palette) + 
  coord_flip() +
  theme_void() +
  theme(legend.position = "none") +
  scale_x_continuous(limits = c(15, 30)) +
  geom_hline(yintercept = 0, colour = "white", size = 1) +
  annotate(geom = "text", y = 0.4, x = 29, label = "Adult\ndistribution", 
           color = "black", size = 4, fontface = 'italic')

# draw the chick weight plot
chick_weight_plot <- 
  ggplot() +
  geom_line(data = chick_growth_results, aes(y = fitm33_w, x = age, 
                                      color = sex),
            size = 1) +
  geom_ribbon(data = chick_growth_results, aes(ymin = uprm33_w, ymax = lwrm33_w, x = age, 
                                        fill = sex), alpha = 0.3) +
  geom_point(data = chick_growth_results, aes(y = weight, x = age, 
                                       fill = sex, 
                                       color = sex),
             alpha = 0.3, size = 2) +
  ylab("Body mass ± 95% CI (g)") +
  xlab("Days since hatching") +
  scale_color_manual(values = sex_palette,
                     name = "Sex",
                     labels = c("Female", "Male")) +
  scale_fill_manual(values = sex_palette,
                    name = "Sex",
                    labels = c("Female", "Male")) +
  theme(legend.position = "none",
        axis.title.x = element_text(size = 16),
        axis.text.x  = element_text(size = 12, angle = 0, vjust = 1, hjust = 0.5)) +
  scale_y_continuous(limits = c(0, 45))

# draw the adult weight plot
adult_weight_plot <- 
  ggplot(adult_measurements, aes(weight)) + 
  geom_density(aes(fill = sex), alpha = 0.3, color = "grey40") + 
  scale_fill_manual(values = sex_palette) + 
  coord_flip() +
  theme_void() +
  theme(legend.position = "none") +
  scale_x_continuous(limits = c(0, 45)) +
  geom_hline(yintercept = 0, colour = "white", size = 1)

# arrange all plots together on one canvas
gridExtra::grid.arrange(chick_tarsus_plot, adult_tarsus_plot, 
                        chick_weight_plot, adult_weight_plot, 
                        ncol = 2, nrow = 2, widths=c(4, 1), heights=c(4, 4))
```

##### Housekeeping

Disconnect from the database

```
dbDisconnect(Ceuta_OPEN)
```

Display R version and package information for time-dependent reproducibility

```
sessionInfo()
```

```
## R version 3.6.1 (2019-07-05)
## Platform: x86_64-apple-darwin15.6.0 (64-bit)
## Running under: macOS Mojave 10.14.5
## 
## Matrix products: default
## BLAS:   /Library/Frameworks/R.framework/Versions/3.6/Resources/lib/libRblas.0.dylib
## LAPACK: /Library/Frameworks/R.framework/Versions/3.6/Resources/lib/libRlapack.dylib
## 
## locale:
## [1] en_US.UTF-8/en_US.UTF-8/en_US.UTF-8/C/en_US.UTF-8/en_US.UTF-8
## 
## attached base packages:
## [1] stats     graphics  grDevices utils     datasets  methods   base     
## 
## other attached packages:
##  [1] plotly_4.9.0       kableExtra_1.1.0   ggplot2_3.2.1     
##  [4] leaflet_2.0.2      webshot_0.5.1      RColorBrewer_1.1-2
##  [7] mapview_2.7.0      rgdal_1.4-4        sp_1.3-1          
## [10] stringr_1.4.0      knitr_1.24         dbplyr_1.4.2      
## [13] dplyr_0.8.3        RSQLite_2.1.2     
## 
## loaded via a namespace (and not attached):
##  [1] nlme_3.1-141       sf_0.8-0           satellite_1.0.1   
##  [4] bit64_0.9-7        httr_1.4.1         rprojroot_1.3-2   
##  [7] tools_3.6.1        backports_1.1.5    utf8_1.1.4        
## [10] R6_2.4.0           KernSmooth_2.23-15 DBI_1.0.0         
## [13] lazyeval_0.2.2     mgcv_1.8-28        colorspace_1.4-1  
## [16] raster_3.0-2       withr_2.1.2        gridExtra_2.3     
## [19] tidyselect_0.2.5   bit_1.1-14         compiler_3.6.1    
## [22] leafem_0.0.1       cli_1.1.0          rvest_0.3.4       
## [25] xml2_1.2.2         labeling_0.3       scales_1.0.0      
## [28] mvtnorm_1.0-11     classInt_0.4-2     readr_1.3.1       
## [31] digest_0.6.21      rmarkdown_1.15     base64enc_0.1-3   
## [34] pkgconfig_2.0.3    htmltools_0.3.6    highr_0.8         
## [37] htmlwidgets_1.3    rlang_0.4.0        rstudioapi_0.10   
## [40] shiny_1.3.2        jsonlite_1.6       crosstalk_1.0.0   
## [43] magrittr_1.5       Formula_1.2-3      Matrix_1.2-17     
## [46] Rcpp_1.0.2         munsell_0.5.0      fansi_0.4.0       
## [49] lifecycle_0.1.0    stringi_1.4.3      yaml_2.2.0        
## [52] grid_3.6.1         blob_1.2.0         parallel_3.6.1    
## [55] promises_1.0.1     crayon_1.3.4       lattice_0.20-38   
## [58] splines_3.6.1      hms_0.5.0          leafpop_0.0.1     
## [61] zeallot_0.1.0      pillar_1.4.2       codetools_0.2-16  
## [64] stats4_3.6.1       glue_1.3.1         evaluate_0.14     
## [67] MBA_0.0-9          data.table_1.12.2  png_0.1-7         
## [70] vctrs_0.2.0        httpuv_1.5.1       gtable_0.3.0      
## [73] purrr_0.3.3        tidyr_1.0.0        assertthat_0.2.1  
## [76] xfun_0.9           mime_0.7           xtable_1.8-4      
## [79] e1071_1.7-2        coda_0.19-3        later_0.8.0       
## [82] survival_2.44-1.1  class_7.3-15       viridisLite_0.3.0 
## [85] tibble_2.1.3       memoise_1.1.0      units_0.6-5       
## [88] bamlss_1.1-1
```
